# RNA binding protein hnRNP-U is required for physiological hypertrophy of skeletal muscle

**DOI:** 10.1101/2020.02.13.944298

**Authors:** Debalina Bagchi, Benjamin D Mason, Kodilichi Baldino, Bin Li, Eun-Joo Lee, Yuteng Zhang, Linh Khanh Chu, Sherif el Raheb, Indranil Sinha, Ronald L Neppl

**Affiliations:** Department of Orthopaedic Surgery, Brigham and Women’s Hospital, Harvard Medical School; Division of Plastic Surgery, Department of Surgery, Brigham and Women’s Hospital, Harvard Medical School

## Abstract

Skeletal muscle has the remarkable ability to modulate its mass in response to physiological changes associated with nutritional input, functional utilization, systemic disease, and age. A decreased responsiveness to anabolic stimuli is thought to contribute significantly to the loss of skeletal muscle mass and strength associated with sarcopenia, however the molecular mechanisms precipitating this are unclear. The signal transduction pathways that control the relative balance between anabolic and catabolic processes are tightly regulated at the transcriptional and post-transcriptional levels. Alternative splicing produces multiple protein isoforms from a single gene in a cell-type-specific manner and in response to environmental cues. We show that sustained activation of Akt1 in *Hnrnpu* deficient mice leads to premature muscle wasting, in part, through impaired autophagy while providing mechanistic insights into the development of anabolic resistance.

## Introduction

Skeletal muscle is a heterogenous mix myofibers that differ in their metabolic and physiological properties and constitutes 40-50% of total body mass. On the ends of this spectrum are the fast glycolytic myofibers that are metabolically optimized to enhance glucose utilization for the rapid production of ATP necessary for short durations of high intensity physical activity, and the slow oxidative myofibers that are metabolically optimized for the continued aerobic production of ATP necessary for prolonged physical activity. A gradual loss of muscle mass and strength is physiologically normal with advancing age (Goodpaster et al., 2006), however a subset of individuals will develop more rapid losses, which may lead to sarcopenia (Evans, 2010). At the cellular level, it is the fast-glycolytic myofibers that are observed to atrophy at a greater rate than slow oxidative fibers (Ciciliot et al., 2013), thus contributing to the loss of muscle power and strength associated with sarcopenia (Lang et al., 2010).

The maintenance of muscle mass is determined by the delicate balance between anabolic and catabolic processes in response to changes in functional utilization, nutritional input, overall health, and age. IGF-1/PI3K/Akt signaling has emerged as a critical regulator of glycolytic muscle growth and metabolism through its activation of mTOR dependent anabolic processes while simultaneously inhibiting Foxo1/3 dependent catabolic processes (Manning and Toker, 2017). Transgenic overexpression of Akt1 in skeletal muscle promotes muscle hypertrophy (Lai et al., 2004), and its selective expression in glycolytic muscles promotes insulin sensitivity in aged mice that display impaired Akt activation (Akasaki et al., 2014). mTOR signals through two complexes, the mTOR containing complex 1 (mTORC1) which both activates anabolic processes while inhibiting autophagy, and the mTOR containing complex 2 (mTORC2) which is the primary Akt1 Ser473 kinase (Manning and Toker, 2017). Though the molecular regulation is complex, changes in energy production and nutrient sensing/storage modulate Akt/mTOR signaling (Manning and Toker, 2017; Yuan et al., 2013), and likely contribute to aged muscles impaired responsiveness to anabolic signals (Breen and Phillips, 2011; Cuthbertson et al., 2005).

Alternative splicing plays a crucial role in the post-transcriptional regulation of gene and protein isoform expression (Lee and Rio, 2015), and is differentially regulated in response to cellular stress (Biamonti and Caceres, 2009; Dutertre et al., 2011). Brain, heart, and skeletal muscle have the highest levels of evolutionary conserved alternatively spliced transcripts (Merkin et al., 2012) and are uniquely sensitive to the mis-splicing of genes (Douglas and Wood, 2013; Scotti and Swanson, 2016). Dysregulated splicing is increasingly recognized as a causal factor promoting cellular dysfunction through changes in the binding properties, enzymatic activities, and intracellular localization of a large percentage of the proteome (Deschenes and Chabot, 2017; Harries et al., 2011; Wang et al., 2018). Age dependent changes in the expression of spliceosome associated factors are implicated as a causal mechanism of cellular aging in the brain through its impact on both metabolism and DNA repair (Mazin et al., 2013; Tollervey et al., 2011). However, the impact of changes in spliceosome associated factor expression on skeletal muscle physiology as a consequence of aging is largely unexplored. Heterogeneous nuclear ribonucleoproteins (hnRNPs) are a large family of RNA-binding proteins (RBPs) involved in critical aspects of nucleic acid metabolism including mRNA stabilization and splicing regulation (Geuens et al., 2016; Martinez-Contreras et al., 2007). Here, we identify hnRNP family member U as being essential for skeletal muscle hypertrophic growth, in part, through modulating the balance between Akt1-dependent anabolic signaling and the energy and nutrient availability to support hypertrophic growth. hnRNP-U expression is shown to be enriched in glycolytic muscles, repressed with age, and necessary for the proper expression and splicing of genes critical for metabolic and growth processes.

## Results

The hnRNP-U protein is expressed in nearly all tissues examined in the adult mouse, with greater expression in the gastrocnemius than in the soleus (**Figure 1A**). However, the expression of hnRNP-U in skeletal muscle decreases abruptly in aged mice (**Figure 1B**). To study the role of hnRNP-U in skeletal muscle growth and function, we crossed Hnrnpu^fl/fl^ mice (Ye et al., 2015) with a mouse line expressing the Cre recombinase under the control of the human skeletal actin promoter (HSA-Cre). HSA-Cre expression starts at E8.5 and continues into adulthood (Miniou et al., 1999). Consistent with a prior report in which Hnrnpu was deleted in both cardiac and skeletal muscle during development (Ye et al., 2015), Hnrnpu^fl/fl^;HSA-Cre (mKO) mice are born in normal Mendelian ratios with no observable abnormalities (**Supplemental Figure 1**). However, Hnrnpu mKO mice continue to develop normally past postnatal day 14 (P14). Laminin stained sections of P21 *Hnrnpu* mutant gastrocnemius (**Figure 1C**) and soleus (**Figure 1F**) are morphologically indistinguishable from littermate controls. Though subtle differences are observed in the myofiber size distribution curves of the gastrocnemius (**Figure 1D**) and soleus (**Figure 1G**), there are no statistically significant differences in the average myofiber cross sectional areas (**Figure 1E and 1H**). Further, we observe no differences in the morphology, myofiber size distribution, or the average myofiber cross sectional area of the tibialis anterior of Hnrnpu mKO mice at P21 (**Supplemental Figure 2**). Collectively, these data indicate embryonic deletion of *Hnrnpu* does not measurably impact the development or post-natal growth of skeletal muscle up to ~P21.

**Figure 1:**
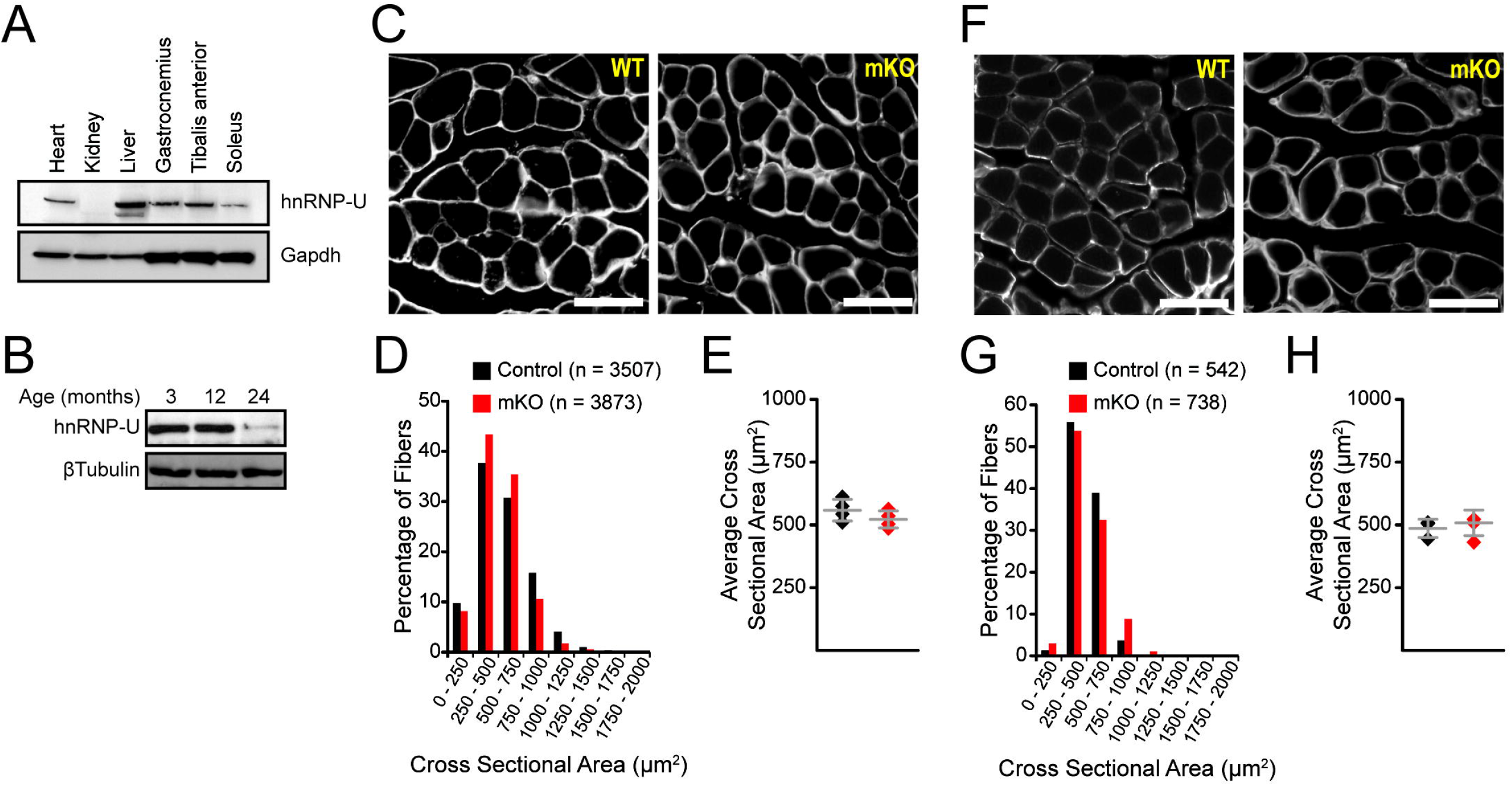
Hnrnpu mKO mice are born phenotypically normal. (A) Representative Western blot of hnRNP-U expression in adult mouse tissues. (B) Representative Western blot depicting hnRNP-U expression in the gastrocnemius with advancing age. (C – H) Data are from n = 3 control and n = 3 mKO mice at P21. (C – E) Immunohistological analyses of gastrocnemius muscle. (F – H) Immunohistological analyses of soleus muscle. (C, F) Laminin staining of muscle transverse sections. The scale bar is 50 μm. (D, G) Myofiber size distribution curves. (E, H) Average myofiber cross sectional areas. Data are mean ± standard deviation. Significance (*), was set at p<0.05.

To explore the role of *Hnrnpu* on skeletal muscle and whole-body growth, we performed longitudinal body weight analyses starting at P21. Consistent with our analyses of mice at P21, male and female mKO mice are indistinguishable by weight from littermate controls up to approximately 4 weeks of age (**Figure 2A**). Growth curve analyses indicate that Hnrnpu mKO mice reach a body weight plateau at approximately 11 – 13 weeks of age, with statistically significant reductions in the body weights of males and females by 6- and 12-weeks of age, respectively. By 7 months of age *Hnrnpu* mutant mice are 100% penetrant for kyphosis, with female mice showing a more severe phenotype by 5-months of age than males by 7-months (**Supplemental Figure 3**). Functional assessments at 3-months of age indicate that deletion of *Hnrnpu* leads to impairments in grip strength (**Figure 2B**) and maximal running speed (**Figure 2C**). Consistent with these analyses, we observe significant reductions in the normalized muscle mass of mKO mice relative to littermate controls (**Figure 2D**). To test the hypothesis that glycolytic muscles are more sensitive to changes in hnRNP-U expression than oxidative muscles, we performed quantitative analyses of myofiber cross sectional areas. Immunofluorescence labeling of laminin and quantification of gastrocnemius myofiber cross sectional areas indicate myofiber atrophy in mKO mice, as indicated by an elevated leftward shift of the myofiber size distribution curve (**Figure 2E and 2F**). Quantitatively, data indicate a significant 21.2% decrease (1479 ± 29 vs. 1878 ± 150 μm^2^, p = 0.011, mKO vs. control) in the average gastrocnemius myofiber cross-sectional area of individual mice (**Figure 2G**). In contrast, the tibialis anterior has only a subtle leftward shift of the myofiber size distribution curve (**Figure 2H and 2I**), whereas the soleus is virtually indistinguishable (**Figure 2K and 2L**) between mKO and littermate controls. While we observe decreases of 7.9% (1541 ± 76 vs. 1674 ± 114 μm^2^, p = 0.058, mKO vs. control) and 3.3% (725 ± 94 vs. 701 ± 98 μm^2^, p = 0.537, mKO vs. control) in the tibialis anterior (**Figure 2J**) and soleus (**Figure 2M**), respectively, these differences fail to reach statistical significance. Consistent with these observations, we observe significant differences in body composition by 3-months of age, notably lean muscle mass, in mKO vs littermate controls (**Table 1**). Collectively, these data indicate that muscles enriched in fast-glycolytic myofibers are particularly sensitive to changes in the hnRNP-U expression necessary for normal skeletal muscle mass, functionality, and strength.

**Figure 2:**
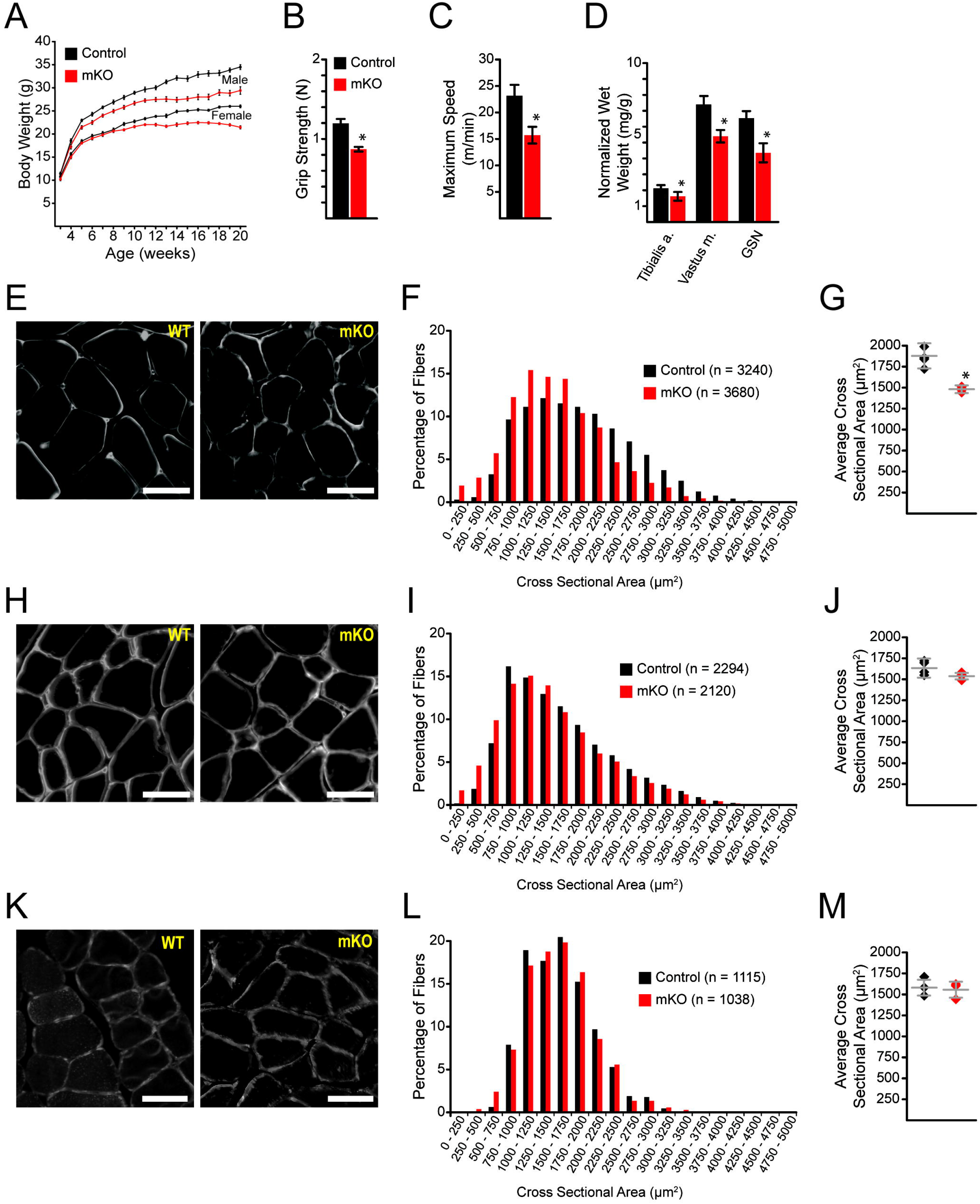
Hnrnpu deletion negatively impacts body growth and muscle function. (A) Growth curves of male and female mice. Data are mean ± SEM from 25 – 40 mice per genotype, depending on age and gender. (B) Grip strength. (C) Maximum running speed. (D) Normalized muscle wet weight. (B – D) Data are mean ± standard deviation from n = 8 control and n = 8 mKO mice. Significance (*) was set a p<0.05. (E - G) Immunohistological analyses of gastrocnemius muscle. (H – J) Immunohistological analyses of tibialis anterior muscle. (K – M) Immunohistological analyses of soleus muscle. (E, H, K) Laminin staining of muscle transverse sections. The scale bar is 50 μm. (F, I, L) Myofiber size distribution curves. (G, J, M) Average myofiber cross sectional areas. Data are mean ± standard deviation. Significance (*), was set at p<0.05. (E – M) Data are from n = 4 control and n = 4 mKO mice. (B – M) Analyses were performed on mice at 3-months of age.

**Table 1:** Morphometric and body composition parameters of WT and hnRNP-U mKO mice at 3 months of age.

|  | **WT (n = 7)** | **cKO (n = 7)** | **p-value** |
| --- | --- | --- | --- |
| Body weight (g) | 28.61 ± 1.46 | 25.70 ± 1.39 | 0.014 |
| Fat (g) | 3.45 ± 0.52 | 3.44 ± 0.57 | 0.995 |
| Lean (g) | 24.26 ± 1.50 | 21.33 ± 1.51 | 0.019 |
| Free water (g) | 0.17 ± 0.10 | 0.14 ± 0.08 | 0.604 |
| Total water (g) | 18.31 ± 1.03 | 14.94 ± 1.14 | 0.003 |

*Data are means ± standard deviation.*

To further explore the biological role of Hnrnpu in skeletal muscle physiology, we assessed its biological role in skeletal muscle regeneration by performing the cryoinjury model in *Hnrnpu* mutant mice at 3-months of age. Morphometric analysis of WT control mice 10-days post injury (DPI) depict robust regeneration characterized by large well-organized myofibers. In contrast, mKO mice exhibit an impaired regenerative response as indicated by numerous small poorly-organized myofibers (**Figure 3A**). Quantitative analyses of histological sections indicate a leftward shift in the myofiber size distribution curve of newly regenerated myofibers in mKO mice relative to controls (**Figure 3B**). To explore whether the age-associated loss of hnRNP-U (**Figure 1B**) contributes to the regenerative deficits of aged muscle, we performed the cryoinjury model on young (3 months) and old (24-26 months) C57BL/6 WT mice. Morphometric analyses of H&E stained sections obtained from young mice exhibit a robust regenerative response at 10 DPI as evidenced by large well-organized regenerated myofibers, whereas old mice exhibit a weak regenerative response characterized by small, poorly-organized myofibers (**Figure 3C**). Quantitative analyses of histological sections indicate a leftward shift in the myofiber size distribution curve of newly regenerated myofibers in aged mice relative to young controls (**Figure 3D**). Overall, the newly regenerated myofibers of young mKO mice are 32% smaller (903 ± 261 vs. 1339 ± 174, mKO vs control) than littermate controls, while the newly regenerated myofibers of old C57BL/6 WT mice are 35% smaller (833 ± 44 vs. 1298 ± 168 μm^2^, old vs. young) than young C57BL/6 controls (**Figure 3E**). Collectively, these observations strongly suggest that the age-associated loss of hnRNP-U contributes to the age-associated reductions in skeletal muscle regeneration.

**Figure 3:**
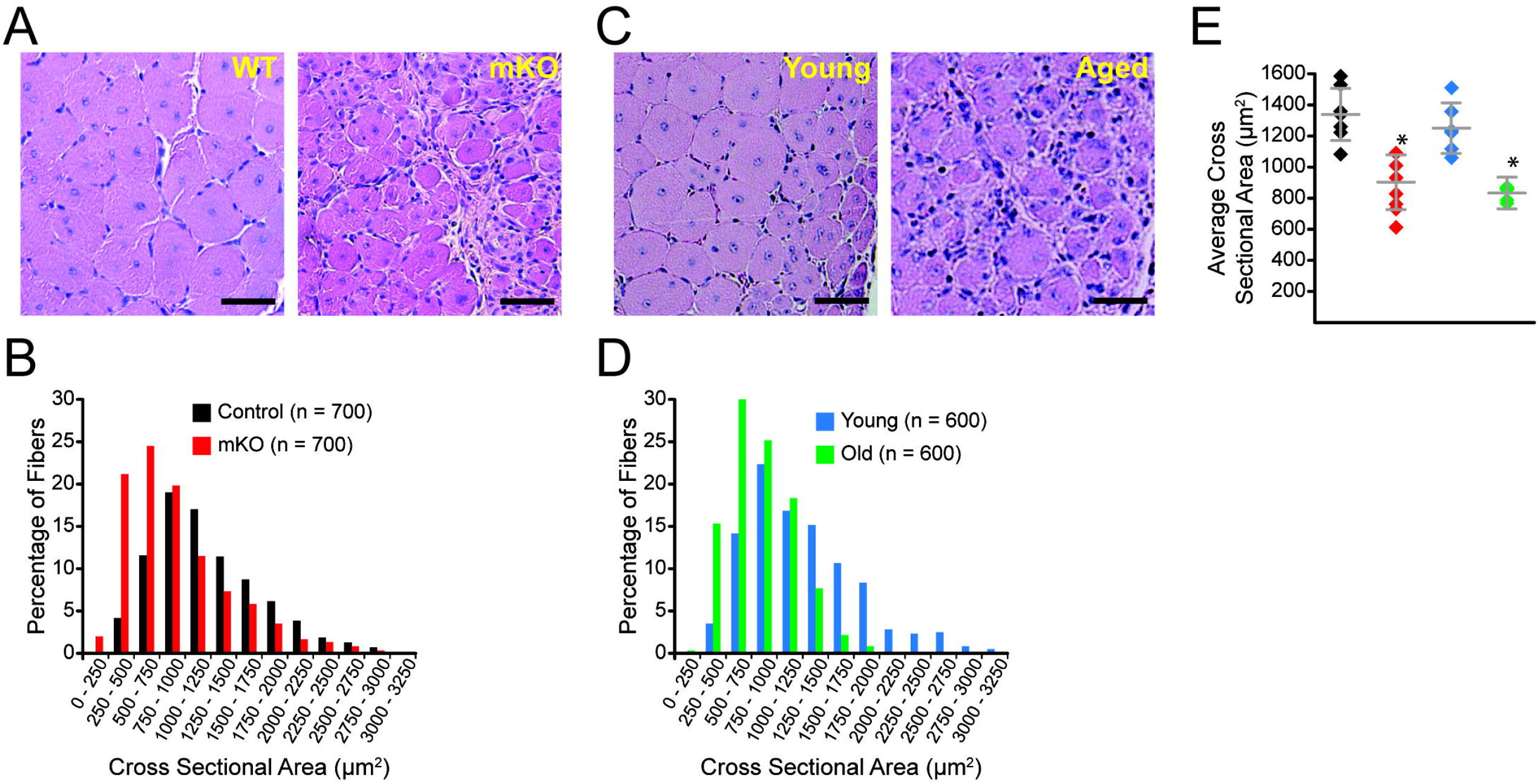
Young *Hnrnpu* mKO mice phenocopy the regenerative response of old mice. (A) H&E stained transverse sections of young (3 months) control and mKO tibialis anterior muscle 10 days post injury. The scale bar is 100 μm. (B) Myofiber size distribution curve of regenerating myofibers from A. (C) H&E stained transverse sections of young (3 months) and old (26 months) tibialis anterior muscles 10 days post injury. The scale bar is 100 μm. (D) Myofiber size distribution curve of regenerating myofibers from C. (E) Dot plot of average regenerated myofiber cross sectional area from n = 7 young *Hnrnpu* control (black diamond), n = 7 young *Hnrnpu* mKO (red diamond), n = 6 young C57BL/6 (blue diamond), and n = 6 old C57BL/6 mice (green diamond). Bars are mean ± standard deviation. Significance (*), was set at p<0.05.

To better understand the molecular processes underlying our observations, we performed whole transcriptome gene and splice variant expression analyses on the gastrocnemius muscle of mice at 3-months of age. We obtained >60 million 150 bp paired end reads per sample with at least 91% of reads mapping to the mouse genome. Differential gene expression (log2FoldChange ± 1.0, p < 0.01) analysis indicate 884 and 1395 genes are increased and decreased in expression (mKO vs. control), respectively (**Figure 4A**). Quantitative PCR validation of RNAseq analyses (data not shown) indicate a marked reduction in the expression of Myostatin, as well as a general repression of the E3 ubiquitin ligases and accessory proteins known to promote muscle atrophy (Bodine et al., 2001; Nowak et al., 2019; Sartori et al., 2013). In addition, genes associated with increased energy expenditures and metabolic stress, including *Sln, Asns, Fgf21, Gdf15, Psat1,* and *Mthfd2* (Balasubramanian et al., 2013; Chung et al., 2017; Fisher and Maratos-Flier, 2016; Maurya et al., 2018; Nilsson et al., 2014 are robustly induced in the muscles of *Hnrnpu* mutants (**Figure 4B**). Interestingly, the induction of lncRNAs appears to portend muscle atrophy and/or metabolic dysfunction, as we observe >3-fold more lncRNAs to be induced than repressed in their expression (Log2FoldChange >2, p < 0.01). To better understand the cellular and molecular processes impacted by the changes in gene expression we performed Gene Ontology enrichment analyses. Our analyses indicate that up-regulated genes are highly associated with inflammatory signaling (**Supplemental Figure 4A**), whereas downregulated genes are associated with energy production and metabolic processes (**Supplemental Figure 4B**). Specifically, we note the marked repression of *Mss51* (Zmynd17) which acts as an inhibitor of oxidative metabolism (Rovira Gonzalez et al., 2019), multiple solute transporters with roles in amino acid and anion/cation (*Slc15a5, Slc30a2, Slc40a1, Slc4a10*) homeostasis (Lin et al., 2015) as well as multiple enzymes involved in the biogenesis of triacylglycerol (*Dgat2l6*) (Yen et al., 2008), amino acids (*Gadl1, Padi2, Amd1/2),* and glycogen (*Gys2*) (**Figure 4C**). The top induced and repressed genes in the gastrocnemius of mKO mice are listed in **Supplemental Data 1**.

**Figure 4:**
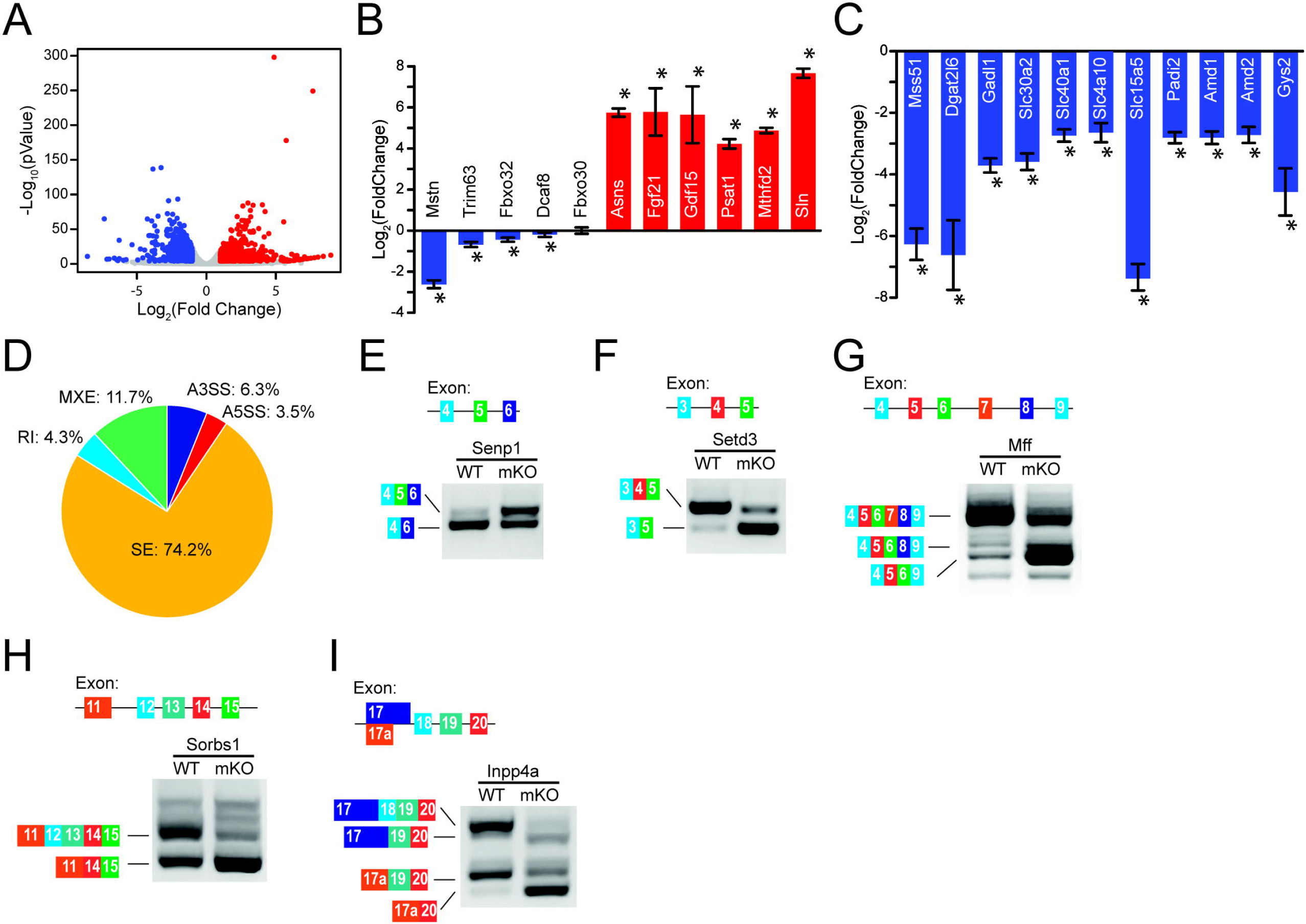
Gene expression and alternative splicing are dysregulated in *Hnrnpu* mKO mice. (A) Whole genome expression profile of *Hnrnpu* mutant gastrocnemius. Red and blue dots depict genes whose expression has increased and decreased (mKO vs control), respectively. Repressed (blue) and induced (red) genes in *Hnrnpu* mutant gastrocnemius associated with (B) muscle atrophy and metabolic stress, and (C) energy production and nutrient transport. (B – C) Data are Log_2_(FoldChange) ± standard error. Significance (*) was set at p<0.0001. (D) Summary of significant differential splicing events between genotypes. |ΔPSI| > 0.1, p < 0.01. (E – I) Examples of differential splicing regulated by hnRNP-U: *Senp1* (E), *Setd3* (F), *Mff* (G), *Sorbs1* (H), and *Inpp4a* (I). SE – Skipped Exon, RI – Retained Intron, MXE – Mutually Exclusive Exon, A3SS – Alternative 3’ Splice Site, A5SS – Alternative 5’ Splice site.

We next examined the impact of *Hnrnpu* deletion on alternative splicing from the paired-end RNAseq data. Using the rMATS program for splicing analysis (Shen et al., 2014), we identified 2909 significant alternative splicing events (|ΔPSI| > 0.1, p < 0.01; **Supplemental Data 2**). The distribution of these events, which include skipped exons (SE), retained introns (RI), mutually exclusive exons (MXE), and the use of alternative 5’ (A5SS) and 3’ (A3SS) splice sites, is shown in **Figure 4D**. These data show that loss of hnRNP-U leads to extensive dysregulation of exon utilization in which SE (74.2%) and MXE (11.7%) events far exceed the utilization of alternative exons (A5SS and A3SS), suggesting that hnRNP-U may be involved in defining intron/exon boundaries. This is in contrast with a prior study in the developing heart, in which RI events were the largest share of alternative splicing events following the loss of hnRNP-U (Ye et al., 2015), thus suggesting that the role of hnRNP-U in pre-mRNA splicing modulation is differentially regulated in a cell type and context-dependent manner.

A more detailed examination of altered splicing events revealed that hnRNP-U is necessary for normal exon utilization in multiple genes, including *Senp1* (**Figure 4E**), *Setd3* (**Figure 4F**), *Mff* (**Figure 4G**), *Sorbs1* (**Figure 4H**), *Inpp4a* (**Figure 4I**). The Sentrin/SUMO-specific protease 1 (*Senp1*) has been previously shown to be involved in regulating mitochondrial biogenesis through enhancing PGC-1α activity (Cai et al., 2012), while mitochondrial fission factor (*Mff*) is a critical regulator of mitochondrial dynamics (Otera et al., 2010). Expression of SET domain containing methyltransferase 3 (*Setd3*) is positively correlated with skeletal muscle hypertrophy (Seaborne et al., 2018) and involved in muscle differentiation via promoting the expression of the myogenic regulatory factors MyoG and Myf6 (Eom et al., 2011). Sorbin and SH3 domain containing protein 1 (*Sorbs1*) is required for PI3K-independent, insulin stimulated glucose uptake (Baumann et al., 2000), whereas the inositol polyphosphate-4-phosphatase 1A (*Inpp4a*) is a PIP2 phosphatase impairing Akt activation (Ivetac et al., 2009). Consistent with these observations, Gene Ontology enrichment analyses indicate that genes with the differential inclusion/exclusion of exons are highly associated with metabolic and signal transduction processes (**Supplemental Figure 4C**). Collectively, these data indicate that *Hnrnpu* is required for the expression and splicing of genes involved in the signal transduction and metabolic processes necessary for normal skeletal muscle growth.

Atrophying muscles, such as occurring with age or age-related sarcopenia, are associated with dysregulated metabolic and signal transduction processes, including that of IGF-1/PI3K/Akt signaling (Karakelides and Nair, 2005; Milman et al., 2016). To explore the impact of *Hnrnpu* deletion on IGF-1/PI3K/Akt signaling, we injected either saline or recombinant IGF-1 into the gastrocnemius muscles of control and mutant mice at ~4 months of age. Quantitative western blot analyses indicate that IGF-1 receptor (IGF-1R) autophosphorylation in response to IGF-1 administration are similar between *Hnrnpu* mutant and control mice (**Figure 5A and 5B**), suggesting that initiation of the IGF-1/PI3K/Akt1 signaling cascade is intact in the absence of Hnrnpu. Upon examination of Akt1 phospho-activation, we observe that IGF-1 stimulation induces a robust induction in Akt1 phosphorylation at Ser473 in control mice (**Figure 5A and 5C**), indicating that IGF-1 stimulation results in the phospho-activation of Akt1. In contrast, we observe that the level of Akt1 Ser473 phosphorylation in vehicle treated *Hnrnpu* mutant mice is comparable to that of IGF-1 stimulated controls (**Figure 5A**). In addition, IGF-1 stimulation is unable to induce further Akt1 Ser473 phosphorylation (**Figure 5A)** suggesting that Akt1 is maximumly activated under normal physiological conditions at this timepoint. Quantification of the phospho-Ser473 to total Akt1 ratio confirms this observation (**Figure 5C**), although the decreased magnitude of the phospho:total Akt1 ratio relative to IGF-1 stimulated controls is due to the increased expression of Akt1 in *Hnrnpu* mutant mice (**Figure 5D**). Collectively, these observations indicate that, in the absence of *Hnrnpu,* Akt1 becomes constitutively activated, and thus desensitized to IGF-1/PI3Ksignaling.

**Figure 5:**
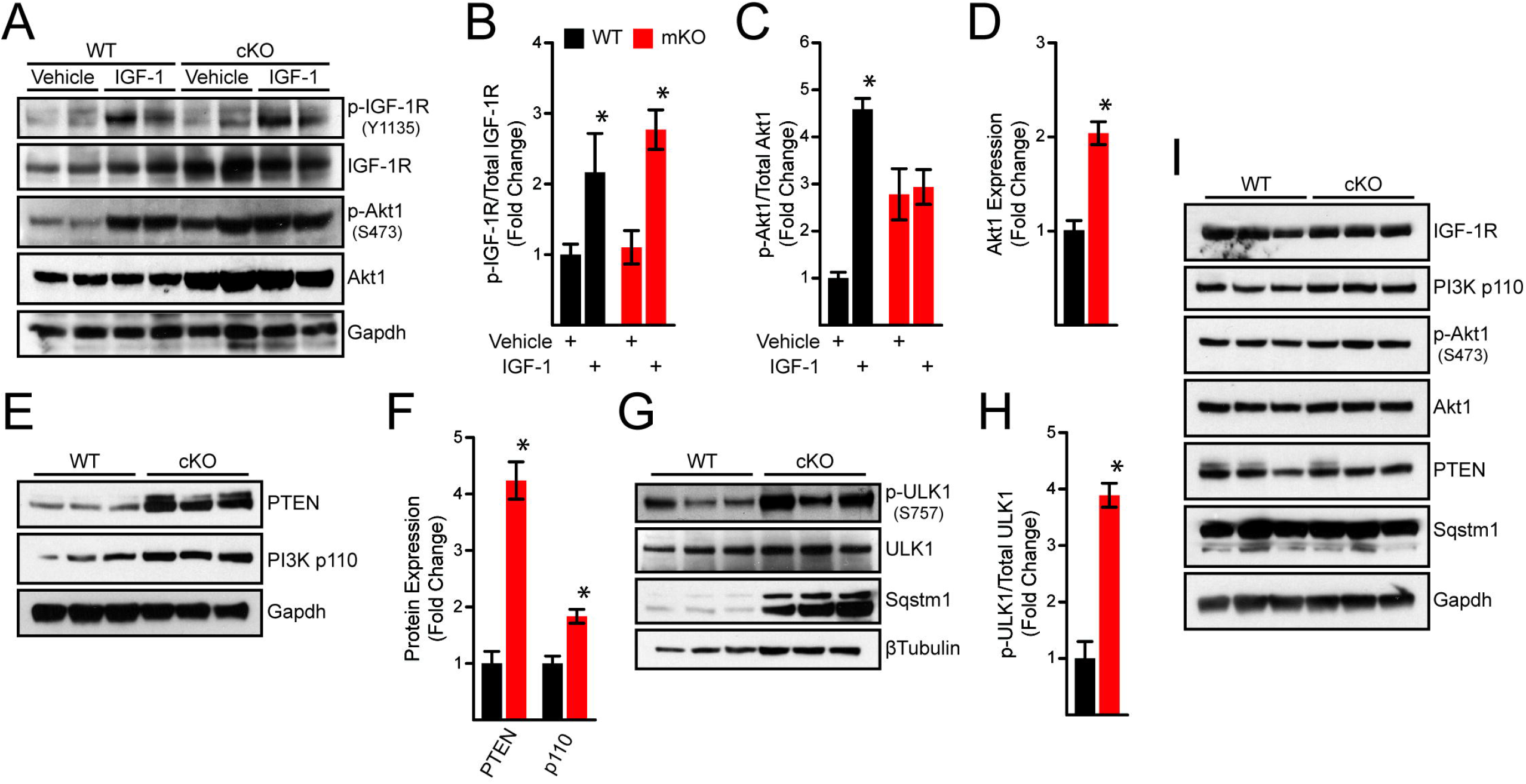
Constitutive activation of Akt1 and impaired autophagic flux in *Hnrnpu* mKO mice. (A) Representative Western blots of gastrocnemius muscle lysates 1 hour following injection of IGF-1. (B – D) Quantification of *A*. Data are mean ± standard deviation from n = 3 independent experiments and are normalized to WT vehicle controls. Significance (*) was set at p<0.05. (E, G) Representative Western blots of unstimulated gastrocnemius muscles at 3 months of age. (F, H) Quantification of *E* and *G*, respectively. (I) Representative Western blots of unstimulated gastrocnemius muscles at P21.

Following up on these observations, we asked whether the signaling downstream of the IGF-1R was biased towards Akt1 activation. First, we examined the expression of the PI3K p110 catalytic subunit. Quantitative western blot analyses indicate that of *Hnrnpu* mutant mice express ~70% more p110 than WT controls (**Figure 5E and 5F**). We next examined PTEN, as it is well established to oppose the Akt1 activating actions of PI3K (Laplante and Sabatini, 2012; Saxton and Sabatini, 2017). Counterintuitively, we observe the expression of PTEN to be increased >4-fold in the gastrocnemius of *Hnrnpu* mutant mice as compared to controls (**Figure 5E and 5F**). However, this observation and the accompanying reductions in body weight (**Figure 2A**), are consistent with a prior report indicating that PTEN overexpression induces postnatal growth impairments through increases in energy expenditure (Garcia-Cao et al., 2012). To better understand the linkages between muscle atrophy, the increased expression of metabolic stress response genes, and the downstream effects of constitutively active Akt1, we assayed for autophagic flux. We observe a marked increase in the expression of the autophagy cargo receptor p62/SQSTM1, and a ~4-fold induction of ULK1 phosphorylation at Ser757 (**Figure 5G and 5H**) indicating that autophagic flux is repressed in an mTORC1 dependent mechanism (Kim et al., 2011; Manning and Toker, 2017).

The postnatal growth of skeletal muscle up to ~P21 is primarily due to the satellite cell contributions to myofiber volume and myonuclei content, after which myonuclear and satellite cell content stabilizes, and hypertrophic signaling begins to dominate (White et al., 2010). As such, we sought to determine whether the satellite cell contributions to muscle growth at this timepoint masked PI3K/Akt signaling derangements. Quantitative Western blot analyses of gastrocnemius muscles of control and mKO mice at P21 indicate no significant differences in the expression or activation of IGF-1R, PI3K p110, Akt1, and PTEN (Figure 5I), indicating the IGF-1/PI3K/Akt signaling is unaltered in *Hnrnpu* mutants at this time point. In addition, the p62/SQSTM1 protein expression in WT and *Hnrnpu* mutants at P21 (Figure 5I) is comparable to that of *Hnrnpu* mutants at 3 months of age (Figure 5G and Supplemental Figure 5), suggesting that the increase in autophagic flux from P21 into adulthood contributes to the hypertrophic growth of skeletal muscle (Masiero et al., 2009).

### Discussion

Disruption of pre-mRNA splicing is increasingly being recognized as significant contributor to muscle and age-associated chronic disease (Deschenes and Chabot, 2017; Faustino and Cooper, 2003). hnRNP-U protein expression in skeletal muscle decreases with age, and genetic deletion of *Hnrnpu* specifically in skeletal muscle leads to premature muscle wasting. Our findings indicate that *Hnrnpu* is essential for the normal physiological growth of skeletal muscle, and further suggest that the loss of hnRNP-U in advanced age may contribute to age-associated functional decline of skeletal muscle. The observed reductions in glycolytic muscle mass and myofiber cross sectional area, impaired regenerative response, and anabolic signaling derangements observed in young *Hnrnpu* mutant mice are consistent with the age-associated losses of glycolytic muscle mass (Ciciliot et al., 2013), regenerative potential (Blau et al., 2015), and anabolic responsiveness (Cuthbertson et al., 2005; Fry et al., 2011). These observations are accompanied by the constitutive activation of Akt/mTOR signaling, decreases in autophagic flux, and transcriptome wide changes in the expression and splicing of genes involved in metabolic and nutrient biosynthetic processes.

IGF-1/PI3K/Akt signaling is a key regulator of glycolytic muscle homeostasis, hypertrophic growth, and metabolism through its activation of mTOR dependent anabolic processes while simultaneously inhibiting catabolic processes (Manning and Toker, 2017; Saxton and Sabatini, 2017). Though the molecular regulation is uncertain, a decreased responsiveness to anabolic stimuli, including growth factors, is thought to play a role in promoting sarcopenia (Cuthbertson et al., 2005). In mice, both the autophosphorylation of IGF-1R, and the sensitivity of Akt1 to IGF-1, are decreased with age (Akasaki et al., 2014). Our results in young *Hnrnpu* mutants indicate that while the autophosphorylation of IGF-1R in response to IGF-1 is unaltered, the phospho-activation of Akt1 is desensitized to IGF-1. These observations indicate that hnRNP-U is essential for the normal signaling activities downstream of IGF-1R and suggest an hnRNP-U independent mechanism modulates the age-associated changes in IGF-1R activation. In contrast with 1-year old non-sarcopenic WT mice (Akasaki et al., 2014), we observe Akt1 to be constitutively active under normal physiological conditions in the atrophied muscles of *Hnrnpu* mutants. Further, we observe Akt signaling in *Hnrnpu* mutants to be desensitized to IGF-1 stimulation, supporting the concept that aberrantly enhanced Akt/mTOR signaling may contribute to the reduced anabolic responsiveness of aging muscles.

Mechanistically, our data indicate that the loss of hnRNP-U leads to a state of cellular stress due to insufficient energy and/or nutrient availability to meet the anabolic demand of sustained Akt1 signaling. Short-term activation of Akt1 signaling specifically within glycolytic myofibers induces changes in expression of genes critical for the activity of pathways involved in the production and storage of energy and nutrients (Wu et al., 2017) along with hypertrophic growth (Izumiya et al., 2008; Wu et al., 2017). Autophagy is induced in response to cellular stress to reduce organelles and macromolecules into amino acids, nucleosides, fatty acids and sugars in support of cellular growth and survival (Rabinowitz and White, 2010). Though it is impaired by Akt/mTOR signaling (Manning and Toker, 2017), insufficient and excessive autophagic flux negatively impact muscle (Mammucari et al., 2007; Masiero et al., 2009), suggesting an inflection point between an autophagic flux that supports muscle hypertrophy and that which promotes muscle wasting. The constitutive activation of mTORC1 in *Tsc1* mKO mice leads to impaired autophagy and muscle atrophy (Castets et al., 2013), whereas inhibition of mTORC1 in sarcopenic rats reduces muscle atrophy, in part, through the restoration of autophagic flux (Joseph et al., 2019). Consistent with these studies, we observe constitutive activation of Akt1 signaling, broad changes in the expression and splicing of genes involved in cellular processes including the production and storage of energy and nutrients, and mTORC1 dependent inhibition of autophagic flux in the atrophied muscles of *Hnrnpu* mutants. The increases in *Asns, Fgf21* and *Gdf15* expression are consistent with a state of nutritional and metabolic stress (Broer and Broer, 2017; Fisher and Maratos-Flier, 2016; Patel et al., 2019) whereas the increase in *Mthfd2*, *Psat1* are likely compensatory mechanisms to mitigate mitochondrial and oxidative stress (Nilsson et al., 2014; Tyynismaa et al., 2010; Yang and Vousden, 2016). Collectively, our studies link the function of *Hnrnpu* to the changes in gene expression and signal transduction underlying muscle atrophy, and implicate hnRNP-U mediate splicing as essential for balancing Akt1 anabolic signaling with the energy and nutrients required for hypertrophic growth. Future detailed analyses dissecting the signaling pathways between Akt1, mTORC1, and autophagy may provide novel insights in muscle wasting disorders, including age related sarcopenia.

## Materials and Methods

### Mouse models

Generation of the Hnrnpu floxed allele has been described previously (Ye et al., 2015). Skeletal muscle specific deletion of Hnrnpu was achieved by crossing with the HSA-Cre (Miniou et al., 1999) mouse line, which expresses the Cre recombinase under the control of the human skeletal alpha-actin promoter. Hnrnpu^fl/fl^ and Hnrnpu^fl/fl^;HSA-Cre mice are on a mixed C57BL/6-129S background. Young (2-3 months of age) and old (24-26 months of age) C57BL/6 mice were obtained from the National Institute of Aging. All animal procedures were approved by the Institutional Animal Care and Use Committee at Brigham and Women’s Hospital.

### Cryoinjury model and histological analysis

Cryoinjury model was performed as previously described (Sinha et al., 2017). Briefly, mice were anesthetized with isoflurane and the skin over the tibialis anterior (TA) was depilated and prepped with sterile alcohol. An incision was made to expose the TA, at which point dry ice was applied directly to the muscle for 5 seconds. The incision was sutured closed immediately after injury. Animals were euthanized at 7 – 10 post injury for histological analyses. Ten (10) images spanning the region of regeneration were acquired per sample. The area of 10 myofibers with central nuclei were chosen randomly for measurement using ImageJ; a total of 100 myofibers were quantified per biological sample. Images were minimally manipulated in Photoshop to linearly increase contrast equally across all pixels.

### IGF-1 response

In vivo signaling downstream of IGF-1 was assessed as previously described (Akasaki et al., 2014). Briefly, mice were anesthetized with isoflurane, and the skin over the gastrocnemius was depilated and prepped with sterile alcohol. Mice were injected with either recombinant human IGF-1 (100 μg), or vehicle (sterile water), directly into the gastrocnemius of the right leg using an insulin syringe. Mice were euthanized 1-hour post injection, and the gastrocnemius was excised and snap-frozen. IGF-1 was purchased from Peprotech Inc (Rocky Hill, NJ).

### Immunohistochemistry and histological analyses

Hind limb muscles were excised, fixed in 4% PFA, and either cryoprotected with 30% sucrose and embedded in Tissue-Tek or dehydrated in ethanol and paraffin embedded. Transverse sections were subjected to immunostaining, nuclei counterstain, imaging, and post-processing as previously described (Neppl et al., 2017). Briefly, fixed tissues were permeabilized with 0.05% Triton X-100 for 20 minutes and blocked with 5% normal donkey serum. Primary antibodies were visualized with cyanine Cy3 or Alexa-Flour 594 conjugated antibodies (Jackson ImmunoResearch Laboratories). Images were acquired at room temperature with a Leica DMi8 microscope with either a 10x or 20x PlanFluotar objective. Approximately 10 – 15 images were acquired from each immunofluorescent stained section. Images were minimally manipulated in Photoshop to linearly increase contrast equally across all pixels.

### Western Blot Analysis

Quantitative western blotting was performed, with minor modifications, according to previously described methods (Neppl et al., 2014). Briefly, tissues were homogenized in 20mM Tris HCl pH 7.5, 150mM NaCl, 1% Triton-X, Halt Protease and Phosphatase Inhibitor Cocktails (Thermo Scientific). Lysates were cleared by centrifugation at 10,000 × g for 10 minutes at 4°C. Protein content was quantified by the DC protein assay (Bio-Rad) with known concentrations of BSA as standards. Protein concentrations were equalized by addition of an appropriate volume of lysis buffer. Primary antibodies were detected by goat antimouse or goat anti-rabbit HRP-conjugated secondary antibodies, and visualized with SuperSignal West Pico PLUS Chemiluminescent Substrate (ThermoFisher Scientific). Films were scanned, and protein band intensities were quantified using ImageJ.

### X-Ray Imaging

Mice were anesthetized with isoflurane and imaged with the Bruker In-Vivo Extreme II Optical/X-ray system for imaging.

### RNAseq and postprocessing for differential gene and splicing analysis

Total RNA from WT and cKO (n = 3, each) gastrocnemius muscle was isolated by TRIzol reagent. The sequencing library was prepared with NEBNext Ultra RNA Library Prep Kit from ribosome depleted RNA, and subjected to HiSeq2500 paired-end strand-specific. Sequencing was performed at a read depth of >60 million reads per biological sample. Following QC and trimming of adapters, reads were mapped using STAR aligner software (Dobin et al., 2013), and counted using featureCounts (Liao et al., 2014). Differential gene expression was quantified using DESeq2 (Love et al., 2014), and analysis of RNA splicing was performed using rMATS (Shen et al., 2014).

### Antibodies

Anti-Akt1 (#2938), Anti-phospho-Akt1 Ser473 (#9271), anti-PTEN (#14642), PI3 Kinase p110α (C73F8), Phospho-IGF-I Receptor β (Tyr1135) (DA7A8), IGF-I Receptor β Antibody #3027, SQSTM1/p62 Antibody (#5114) and anti-β-Tubulin (#2146) were purchased from Cell Signaling Technologies. Anti-GAPDH, clone 6C5 was purchased from EMD Millipore.

## Acknowledgments

The authors would like to thank Dr. Tom Maniatis for the Hnrnpu mouse line and Teri Bowman for her technical assistance with histology processing. This work was supported by funds from the BWH Department of Orthopaedic Surgery to RLN and K76AG059996 to IS.

## References

Akasaki, Y., N. Ouchi, Y. Izumiya, B.L. Bernardo, N.K. Lebrasseur, and K. Walsh. 2014. Glycolytic fast-twitch muscle fiber restoration counters adverse age-related changes in body composition and metabolism. Aging Cell. 13:80–91.

Balasubramanian, M.N., E.A. Butterworth, and M.S. Kilberg. 2013. Asparagine synthetase: regulation by cell stress and involvement in tumor biology. Am J Physiol Endocrinol Metab. 304:E789–799.

Baumann, C.A., V. Ribon, M. Kanzaki, D.C. Thurmond, S. Mora, S. Shigematsu, P.E. Bickel, J.E. Pessin, and A.R. Saltiel. 2000. CAP defines a second signalling pathway required for insulin-stimulated glucose transport. Nature. 407:202–207.

Biamonti, G., and J.F. Caceres. 2009. Cellular stress and RNA splicing. Trends Biochem Sci. 34:146–153.

Blau, H.M., B.D. Cosgrove, and A.T. Ho. 2015. The central role of muscle stem cells in regenerative failure with aging. Nat Med. 21:854–862.

Bodine, S.C., E. Latres, S. Baumhueter, V.K. Lai, L. Nunez, B.A. Clarke, W.T. Poueymirou, F.J. Panaro, E. Na, K. Dharmarajan, Z.Q. Pan, D.M. Valenzuela, T.M. DeChiara, T.N. Stitt, G.D. Yancopoulos, and D.J. Glass. 2001. Identification of ubiquitin ligases required for skeletal muscle atrophy. Science. 294:1704–1708.

Breen, L., and S.M. Phillips. 2011. Skeletal muscle protein metabolism in the elderly: Interventions to counteract the ‘anabolic resistance’ of ageing. Nutr Metab (Lond). 8:68.

Broer, S., and A. Broer. 2017. Amino acid homeostasis and signalling in mammalian cells and organisms. Biochem J. 474:1935–1963.

Cai, R., T. Yu, C. Huang, X. Xia, X. Liu, J. Gu, S. Xue, E.T. Yeh, and J. Cheng. 2012. SUMO-specific protease 1 regulates mitochondrial biogenesis through PGC-1alpha. J Biol Chem. 287:44464–44470.

Castets, P., S. Lin, N. Rion, S. Di Fulvio, K. Romanino, M. Guridi, S. Frank, L.A. Tintignac, M. Sinnreich, and M.A. Ruegg. 2013. Sustained activation of mTORC1 in skeletal muscle inhibits constitutive and starvation-induced autophagy and causes a severe, late-onset myopathy. Cell Metab. 17:731–744.

Chung, H.K., D. Ryu, K.S. Kim, J.Y. Chang, Y.K. Kim, H.S. Yi, S.G. Kang, M.J. Choi, S.E. Lee, S.B. Jung, M.J. Ryu, S.J. Kim, G.R. Kweon, H. Kim, J.H. Hwang, C.H. Lee, S.J. Lee, C.E. Wall, M. Downes, R.M. Evans, J. Auwerx, and M. Shong. 2017. Growth differentiation factor 15 is a myomitokine governing systemic energy homeostasis. J Cell Biol. 216:149–165.

Ciciliot, S., A.C. Rossi, K.A. Dyar, B. Blaauw, and S. Schiaffino. 2013. Muscle type and fiber type specificity in muscle wasting. Int J Biochem Cell Biol. 45:2191–2199.

Cuthbertson, D., K. Smith, J. Babraj, G. Leese, T. Waddell, P. Atherton, H. Wackerhage, P.M. Taylor, and M.J. Rennie. 2005. Anabolic signaling deficits underlie amino acid resistance of wasting, aging muscle. FASEB J. 19:422–424.

Deschenes, M., and B. Chabot. 2017. The emerging role of alternative splicing in senescence and aging. Aging Cell. 16:918–933.

Dobin, A., C.A. Davis, F. Schlesinger, J. Drenkow, C. Zaleski, S. Jha, P. Batut, M. Chaisson, and T.R. Gingeras. 2013. STAR: ultrafast universal RNA-seq aligner. Bioinformatics. 29:15–21.

Douglas, A.G., and M.J. Wood. 2013. Splicing therapy for neuromuscular disease. Mol Cell Neurosci. 56:169–185.

Dutertre, M., G. Sanchez, J. Barbier, L. Corcos, and D. Auboeuf. 2011. The emerging role of pre-messenger RNA splicing in stress responses: sending alternative messages and silent messengers. RNA Biol. 8:740–747.

Eom, G.H., K.B. Kim, J.H. Kim, J.Y. Kim, J.R. Kim, H.J. Kee, D.W. Kim, N. Choe, H.J. Park, H.J. Son, S.Y. Choi, H. Kook, and S.B. Seo. 2011. Histone methyltransferase SETD3 regulates muscle differentiation. J Biol Chem. 286:34733–34742.

Evans, W.J. 2010. Skeletal muscle loss: cachexia, sarcopenia, and inactivity. Am J Clin Nutr. 91:1123S–1127S.

Faustino, N.A., and T.A. Cooper. 2003. Pre-mRNA splicing and human disease. Genes Dev. 17:419–437.

Fisher, F.M., and E. Maratos-Flier. 2016. Understanding the Physiology of FGF21. Annu Rev Physiol. 78:223–241.

Fry, C.S., M.J. Drummond, E.L. Glynn, J.M. Dickinson, D.M. Gundermann, K.L. Timmerman, D.K. Walker, S. Dhanani, E. Volpi, and B.B. Rasmussen. 2011. Aging impairs contraction-induced human skeletal muscle mTORC1 signaling and protein synthesis. Skelet Muscle. 1:11.

Garcia-Cao, I., M.S. Song, R.M. Hobbs, G. Laurent, C. Giorgi, V.C. de Boer, D. Anastasiou, K. Ito, A.T. Sasaki, L. Rameh, A. Carracedo, M.G. Vander Heiden, L.C. Cantley, P. Pinton, M.C. Haigis, and P.P. Pandolfi. 2012. Systemic elevation of PTEN induces a tumor-suppressive metabolic state. Cell. 149:49–62.

Geuens, T., D. Bouhy, and V. Timmerman. 2016. The hnRNP family: insights into their role in health and disease. Hum Genet. 135:851–867.

Goodpaster, B.H., S.W. Park, T.B. Harris, S.B. Kritchevsky, M. Nevitt, A.V. Schwartz, E.M. Simonsick, F.A. Tylavsky, M. Visser, and A.B. Newman. 2006. The loss of skeletal muscle strength, mass, and quality in older adults: the health, aging and body composition study. J Gerontol A Biol Sci Med Sci. 61:1059–1064.

Harries, L.W., D. Hernandez, W. Henley, A.R. Wood, A.C. Holly, R.M. Bradley-Smith, H. Yaghootkar, A. Dutta, A. Murray, T.M. Frayling, J.M. Guralnik, S. Bandinelli, A. Singleton, L. Ferrucci, and D. Melzer. 2011. Human aging is characterized by focused changes in gene expression and deregulation of alternative splicing. Aging Cell. 10:868–878.

Ivetac, I., R. Gurung, S. Hakim, K.A. Horan, D.A. Sheffield, L.C. Binge, P.W. Majerus, T. Tiganis, and C.A. Mitchell. 2009. Regulation of PI(3)K/Akt signalling and cellular transformation by inositol polyphosphate 4-phosphatase-1. EMBO Rep. 10:487–493.

Izumiya, Y., T. Hopkins, C. Morris, K. Sato, L. Zeng, J. Viereck, J.A. Hamilton, N. Ouchi, N.K. LeBrasseur, and K. Walsh. 2008. Fast/Glycolytic muscle fiber growth reduces fat mass and improves metabolic parameters in obese mice. Cell Metab. 7:159–172.

Joseph, G.A., S.X. Wang, C.E. Jacobs, W. Zhou, G.C. Kimble, H.W. Tse, J.K. Eash, T. Shavlakadze, and D.J. Glass. 2019. Partial Inhibition of mTORC1 in Aged Rats Counteracts the Decline in Muscle Mass and Reverses Molecular Signaling Associated with Sarcopenia. Mol Cell Biol. 39.

Karakelides, H., and K.S. Nair. 2005. Sarcopenia of aging and its metabolic impact. Curr Top Dev Biol. 68:123–148.

Kim, J., M. Kundu, B. Viollet, and K.L. Guan. 2011. AMPK and mTOR regulate autophagy through direct phosphorylation of Ulk1. Nat Cell Biol. 13:132–141.

Lai, K.M., M. Gonzalez, W.T. Poueymirou, W.O. Kline, E. Na, E. Zlotchenko, T.N. Stitt, A.N. Economides, G.D. Yancopoulos, and D.J. Glass. 2004. Conditional activation of akt in adult skeletal muscle induces rapid hypertrophy. Mol Cell Biol. 24:9295–9304.

Lang, T., T. Streeper, P. Cawthon, K. Baldwin, D.R. Taaffe, and T.B. Harris. 2010. Sarcopenia: etiology, clinical consequences, intervention, and assessment. Osteoporos Int. 21:543–559.

Laplante, M., and D.M. Sabatini. 2012. mTOR signaling in growth control and disease. Cell. 149:274–293.

Lee, Y., and D.C. Rio. 2015. Mechanisms and Regulation of Alternative Pre-mRNA Splicing. Annu Rev Biochem. 84:291–323.

Liao, Y., G.K. Smyth, and W. Shi. 2014. featureCounts: an efficient general purpose program for assigning sequence reads to genomic features. Bioinformatics. 30:923–930.

Lin, L., S.W. Yee, R.B. Kim, and K.M. Giacomini. 2015. SLC transporters as therapeutic targets: emerging opportunities. Nat Rev Drug Discov. 14:543–560.

Love, M.I., W. Huber, and S. Anders. 2014. Moderated estimation of fold change and dispersion for RNA-seq data with DESeq2. Genome Biol. 15:550.

Mammucari, C., G. Milan, V. Romanello, E. Masiero, R. Rudolf, P. Del Piccolo, S.J. Burden, R. Di Lisi, C. Sandri, J. Zhao, A.L. Goldberg, S. Schiaffino, and M. Sandri. 2007. FoxO3 controls autophagy in skeletal muscle in vivo. Cell Metab. 6:458–471.

Manning, B.D., and A. Toker. 2017. AKT/PKB Signaling: Navigating the Network. Cell. 169:381–405.

Martinez-Contreras, R., P. Cloutier, L. Shkreta, J.F. Fisette, T. Revil, and B. Chabot. 2007. hnRNP proteins and splicing control. Adv Exp Med Biol. 623:123–147.

Masiero, E., L. Agatea, C. Mammucari, B. Blaauw, E. Loro, M. Komatsu, D. Metzger, C. Reggiani, S. Schiaffino, and M. Sandri. 2009. Autophagy is required to maintain muscle mass. Cell Metab. 10:507–515.

Maurya, S.K., J.L. Herrera, S.K. Sahoo, F.C.G. Reis, R.B. Vega, D.P. Kelly, and M. Periasamy. 2018. Sarcolipin Signaling Promotes Mitochondrial Biogenesis and Oxidative Metabolism in Skeletal Muscle. Cell Rep. 24:2919–2931.

Mazin, P., J. Xiong, X. Liu, Z. Yan, X. Zhang, M. Li, L. He, M. Somel, Y. Yuan, Y.P. Phoebe Chen, N. Li, Y. Hu, N. Fu, Z. Ning, R. Zeng, H. Yang, W. Chen, M. Gelfand, and P. Khaitovich. 2013. Widespread splicing changes in human brain development and aging. Mol Syst Biol. 9:633.

Merkin, J., C. Russell, P. Chen, and C.B. Burge. 2012. Evolutionary dynamics of gene and isoform regulation in Mammalian tissues. Science. 338:1593–1599.

Milman, S., D.M. Huffman, and N. Barzilai. 2016. The Somatotropic Axis in Human Aging: Framework for the Current State of Knowledge and Future Research. Cell Metab. 23:980–989.

Miniou, P., D. Tiziano, T. Frugier, N. Roblot, M. Le Meur, and J. Melki. 1999. Gene targeting restricted to mouse striated muscle lineage. Nucleic Acids Res. 27:e27.

Neppl, R.L., M. Kataoka, and D.Z. Wang. 2014. Crystallin-alphaB Regulates Skeletal Muscle Homeostasis via Modulation of Argonaute2 Activity. J Biol Chem. 289:17240–17248.

Neppl, R.L., C.L. Wu, and K. Walsh. 2017. lncRNA Chronos is an aging-induced inhibitor of muscle hypertrophy. J Cell Biol. 216:3497–3507.

Nilsson, R., M. Jain, N. Madhusudhan, N.G. Sheppard, L. Strittmatter, C. Kampf, J. Huang, A. Asplund, and V.K. Mootha. 2014. Metabolic enzyme expression highlights a key role for MTHFD2 and the mitochondrial folate pathway in cancer. Nat Commun. 5:3128.

Nowak, M., B. Suenkel, P. Porras, R. Migotti, F. Schmidt, M. Kny, X. Zhu, E.E. Wanker, G. Dittmar, J. Fielitz, and T. Sommer. 2019. DCAF8, a novel MuRF1 interaction partner, promotes muscle atrophy. J Cell Sci. 132.

Otera, H., C. Wang, M.M. Cleland, K. Setoguchi, S. Yokota, R.J. Youle, and K. Mihara. 2010. Mff is an essential factor for mitochondrial recruitment of Drp1 during mitochondrial fission in mammalian cells. J Cell Biol. 191:1141–1158.

Patel, S., A. Alvarez-Guaita, A. Melvin, D. Rimmington, A. Dattilo, E.L. Miedzybrodzka, I. Cimino, A.C. Maurin, G.P. Roberts, C.L. Meek, S. Virtue, L.M. Sparks, S.A. Parsons, L.M. Redman, G.A. Bray, A.P. Liou, R.M. Woods, S.A. Parry, P.B. Jeppesen, A.J. Kolnes, H.P. Harding, D. Ron, A. Vidal-Puig, F. Reimann, F.M. Gribble, C.J. Hulston, I.S. Farooqi, P. Fafournoux, S.R. Smith, J. Jensen, D. Breen, Z. Wu, B.B. Zhang, A.P. Coll, D.B. Savage, and S. O’Rahilly. 2019. GDF15 Provides an Endocrine Signal of Nutritional Stress in Mice and Humans. Cell Metab. 29:707–718 e708.

Rabinowitz, J.D., and E. White. 2010. Autophagy and metabolism. Science. 330:1344–1348.

Rovira Gonzalez, Y.I., A.L. Moyer, N.J. LeTexier, A.D. Bratti, S. Feng, C. Sun, T. Liu, J. Mula, P. Jha, S.R. Iyer, R.M. Lovering, B. O’Rourke, H.L. Noh, S. Suk, J.K. Kim, G.K. Essien Umanah, and K.R. Wagner. 2019. Mss51 deletion enhances muscle metabolism and glucose homeostasis in mice. JCI Insight. 4.

Sartori, R., E. Schirwis, B. Blaauw, S. Bortolanza, J. Zhao, E. Enzo, A. Stantzou, E. Mouisel, L. Toniolo, A. Ferry, S. Stricker, A.L. Goldberg, S. Dupont, S. Piccolo, H. Amthor, and M. Sandri. 2013. BMP signaling controls muscle mass. Nat Genet. 45:1309–1318.

Saxton, R.A., and D.M. Sabatini. 2017. mTOR Signaling in Growth, Metabolism, and Disease. Cell. 168:960–976.

Scotti, M.M., and M.S. Swanson. 2016. RNA mis-splicing in disease. Nat Rev Genet. 17:19–32.

Seaborne, R.A., J. Strauss, M. Cocks, S. Shepherd, T.D. O’Brien, K.A. van Someren, P.G. Bell, C. Murgatroyd, J.P. Morton, C.E. Stewart, and A.P. Sharples. 2018. Human Skeletal Muscle Possesses an Epigenetic Memory of Hypertrophy. Sci Rep. 8:1898.

Shen, S., J.W. Park, Z.X. Lu, L. Lin, M.D. Henry, Y.N. Wu, Q. Zhou, and Y. Xing. 2014. rMATS: robust and flexible detection of differential alternative splicing from replicate RNA-Seq data. Proc Natl Acad Sci U S A. 111:E5593–5601.

Sinha, I., D. Sakthivel, B.A. Olenchock, C.R. Kruse, J. Williams, D.E. Varon, J.D. Smith, A.L. Madenci, K. Nuutila, and A.J. Wagers. 2017. Prolyl Hydroxylase Domain-2 Inhibition Improves Skeletal Muscle Regeneration in a Male Murine Model of Obesity. Front Endocrinol (Lausanne). 8:153.

Tollervey, J.R., Z. Wang, T. Hortobagyi, J.T. Witten, K. Zarnack, M. Kayikci, T.A. Clark, A.C. Schweitzer, G. Rot, T. Curk, B. Zupan, B. Rogelj, C.E. Shaw, and J. Ule. 2011. Analysis of alternative splicing associated with aging and neurodegeneration in the human brain. Genome Res. 21:1572–1582.

Tyynismaa, H., C.J. Carroll, N. Raimundo, S. Ahola-Erkkila, T. Wenz, H. Ruhanen, K. Guse, A. Hemminki, K.E. Peltola-Mjosund, V. Tulkki, M. Oresic, C.T. Moraes, K. Pietilainen, I. Hovatta, and A. Suomalainen. 2010. Mitochondrial myopathy induces a starvation-like response. Hum Mol Genet. 19:3948–3958.

Wang, K., D. Wu, H. Zhang, A. Das, M. Basu, J. Malin, K. Cao, and S. Hannenhalli. 2018. Comprehensive map of age-associated splicing changes across human tissues and their contributions to age-associated diseases. Sci Rep. 8:10929.

White, R.B., A.S. Bierinx, V.F. Gnocchi, and P.S. Zammit. 2010. Dynamics of muscle fibre growth during postnatal mouse development. BMC Dev Biol. 10:21.

Wu, C.L., Y. Satomi, and K. Walsh. 2017. RNA-seq and metabolomic analyses of Akt1-mediated muscle growth reveals regulation of regenerative pathways and changes in the muscle secretome. BMC Genomics. 18:181.

Yang, M., and K.H. Vousden. 2016. Serine and one-carbon metabolism in cancer. Nat Rev Cancer. 16:650–662.

Ye, J., N. Beetz, S. O’Keeffe, J.C. Tapia, L. Macpherson, W.V. Chen, R. Bassel-Duby, E.N. Olson, and T. Maniatis. 2015. hnRNP U protein is required for normal pre-mRNA splicing and postnatal heart development and function. Proc Natl Acad Sci U S A. 112:E3020–3029.

Yen, C.L., S.J. Stone, S. Koliwad, C. Harris, and R.V. Farese, Jr. 2008. Thematic review series: glycerolipids. DGAT enzymes and triacylglycerol biosynthesis. J Lipid Res. 49:2283–2301.

Yuan, H.X., Y. Xiong, and K.L. Guan. 2013. Nutrient sensing, metabolism, and cell growth control. Mol Cell. 49:379–387.

