## Supplemental Figures for "RNA binding protein hnRNP-U is required for physiological hypertrophy of skeletal muscle"

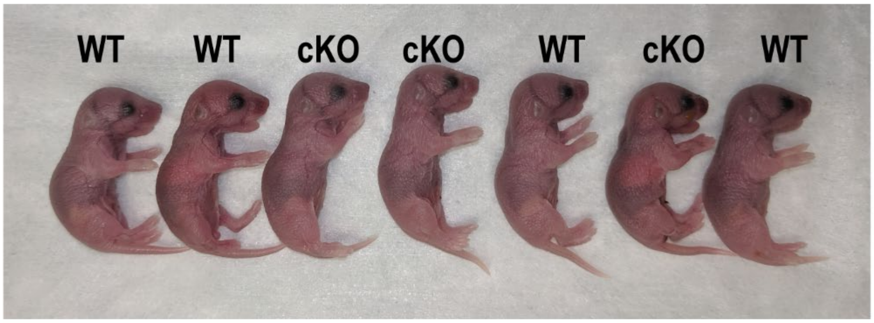


**Supplemental Figure 1: Representative image of WT and Hnrnpu mutant littermates at P0.5.**

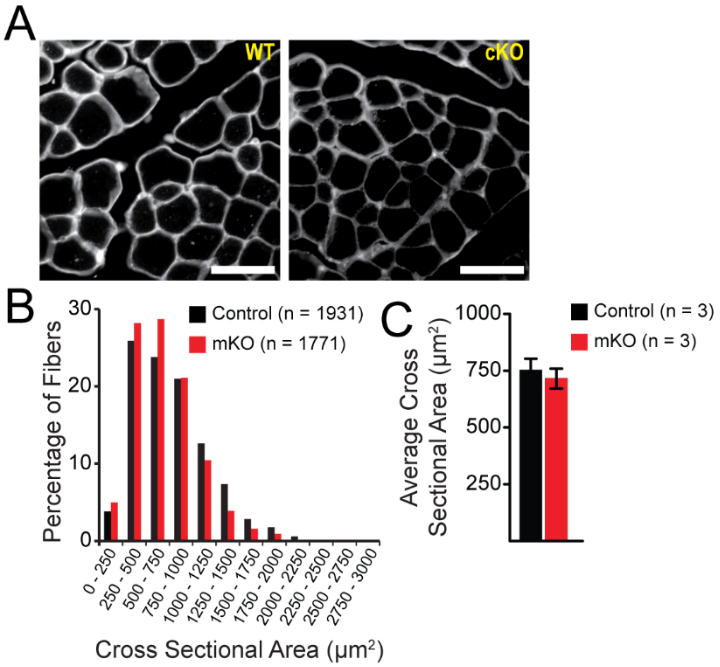


**Supplemental Figure 2: Morphometric analysis of tibialis anterior muscles at P21.** (A) Representative images of laminin stained cross sections obtained from WT and Hnrnpu mutant muscles. Scale bar is 50 µm. (B) Myofiber size distribution. (C) Mean myofiber cross-sectional area across genotypes. Data are mean ± standard deviation.

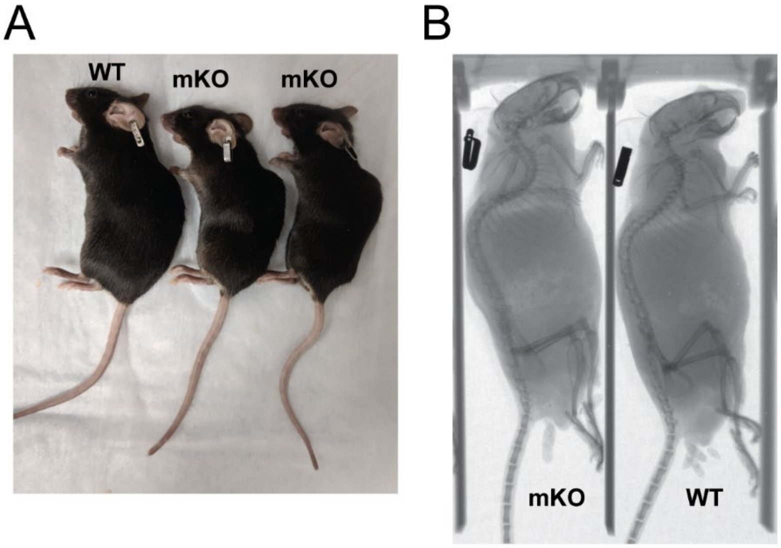


**Supplemental Figure 3: Hnnrpu mKO mice develop kyphosis.** (A) Representative images of female littermates at 5 months of age. (B) Representative X-ray micrograph of male littermates at 6 – 7 months of age.

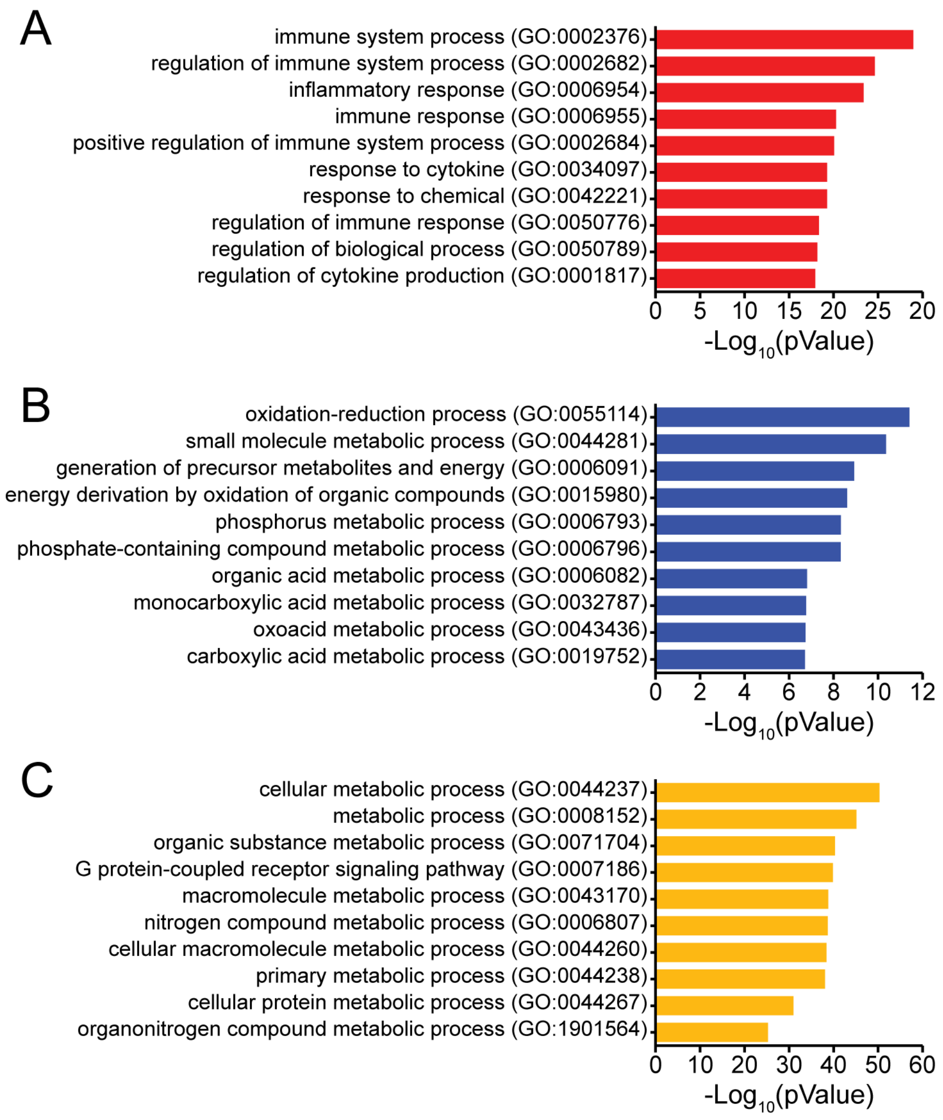


**Supplemental Figure 4: Gene Ontology Gene Enrichment Analysis of RNAseq data.** To GO terms of genes whose expression is increased (A), decreased (B), or differentially spliced (C) in *Hnrnpu* mutants.

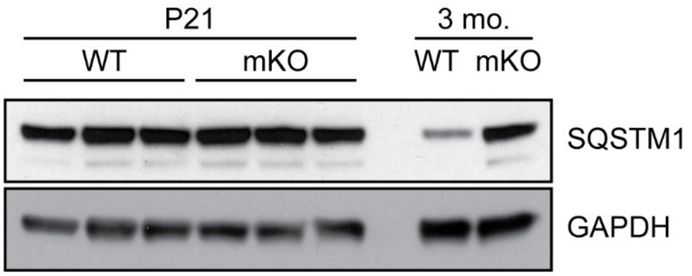


**Supplemental Figure 5: Age associated autophagic flux.** Western blot analysis of gastrocnemius muscles of WT and Hnrnpu mKO mice at P21 and 3 months of age. Equal amounts of total protein (30 µg) were loaded in each lane.
